# Combined oncolytic adenovirus carrying MnSOD and mK5 genes both regulated by survivin promoter has synergistic inhibitory effect on gastric cancer

**DOI:** 10.1101/2021.04.08.438934

**Authors:** Shanshan Liu, JinQing Hu, Jinfa Gu, Aimin Ni, Wenhao Tang, Xinyuan Liu

## Abstract

Gastric cancer (GC) is one of the major causes of cancer related mortality. The use of oncolytic virus for cancer gene-virotherapy is a new approach for the treatment of human cancers. In this study, a novel *Survivin* promoter driven recombinant oncolytic adenovirus carrying *mK5* or *MnSOD* gene was constructed, which was modified after deletion of E1B gene. Human plasminogen Kringle 5 mutant (*mK5*) and manganese superoxide dismutase (*MnSOD*) are both potential tumor suppressor genes. To construct Ad-Surp-*mK5* and Ad-Surp-*MnSOD* oncolytic adenovirus, we hypothesized that the combination of the two viruses would enhance the therapeutic efficacy of GC as compared to the virus alone. The results of the *in vitro* experiments revealed that the combination of adenovirus carrying *mK5* and *MnSOD* gene exhibited stronger cytotoxicity to GC cell lines as compared to the virus alone, Additionally, the virus could selectively kill cancer cells and human somatic cells. Cell staining, flow cytometry and western blot analysis showed that the combination of two adenovirus containing therapeutic genes could promote the apoptosis of cancer cells. *In vivo* experiments further verified that Ad-Surp-*mK5* in combination with Ad-Surp-*MnSOD* exhibited significant inhibitory effect on the growth of GC tumor xenograft as compared to the virus alone, and no significant difference was observed in the body weight of treatment and the normal mice. In conclusion, the combination of our two newly constructed recombinant oncolytic adenovirus containing *mK5* or *MnSOD* therapeutic genes could significantly inhibit gastric cancer growth by inducing apoptosis, suggestive of its potential for GC therapy.

## Introduction

Gastric cancer (GC) is the fifth most prevalent cancer worldwide and the third leading cause of cancer related mortality [1]. GC is one of the most common malignant tumors of the digestive system. At present, surgery is considered as the only radical cure. The 5-year survival rate of early gastric cancer may reach 95%, but due to late diagnosis, most of the patients are diagnosed at advanced stages [2]. Despite advances in surgical techniques, radiotherapy, chemotherapy and neoadjuvant therapy [3], GC still remains a serious global health burden [4]. Many cancers can be cured by surgery, yet most of the world’s population does not have access to safe, affordable and timely cancer surgery [5].

Worldwide interest in oncolytic viruses (OVs), a powerful new oncology drug, has increased remarkably with the approval of the first OV (Talimogene Laherparepvec) by the US Food and Drug Administration [6]. However, after extensive research oncolytic virus therapy is limited in the treatment of solid tumors.We used a novel strategy called targeted gene-virotherapy of cancer (CTGVT) that combined the advantages of gene therapy and oncolytic virus therapy to produce a stronger anti-tumor effect than either gene or oncolytic virus therapy alone [7]. For example, ONYX-015 is an oncolytic adenovirus (Ad) with deletion of its viral protein E1B-55kDa gene. Due to the inability of the mutated protein to degrade P53, the selectivity of tumor replication is enhanced. The early gene E1B gene encoded by Ads mainly contains two kinds of polypeptides E1B-19kDa and E1B-55kDa. E1B-19kDa is a homologous of Bcl-2 which can prevent E1A-induced apoptosis by interfering with Bak, Bax interaction. By E1B-19 kDa protein induced inhibition of FAS mediated apoptosis and inhibition of apoptosis of cancer cells through other apoptotic pathways, Ad knockout E1B-19 kDa after virus replication in restricted in normal cells and cancer cells are not suppressed [8]. In the host,the E1B-55 kDa protein binds to P53 which consequently leads to the inhibition of the viral replication. Therefore, OV-knockout E1B-55 kDa can’t replicate in the normal cells, but the virus replicates normally in tumor cells lacking P53 function and causes tumor cells dissolve. However, the oncolytic adenovirus produced simply by deleting viral genes is not selective enough to prevent unnecessary tissue damage. Recently, tumor-specific promoters have become the most popular tool for controlling the oncolytic adenoviruses that strictly target cancer cells [9]. *Survivin*, a member of the apoptosis-inhibiting protein family, is overexpressed in several types of cancers but is not expressed in differentiated normal tissues [10]. The results suggest that *Survivin* promoter is a specific promoter for a variety of cancers and may play a role in cancer gene therapy, and that *Survivin* promoter is more tumor-specific than CMV promoter *in vivo* [11].

The anti-angiogenic protein K5 (human plasminogen Kringle 5) forms the *mK5* mutation by mutation of 71 leucine residue to arginine [12]. Studies have shown that *mK5* and K5 can considerably inhibit the proliferation of human umbilical vein endothelial cells (HUVECs) and induce the apoptosis of vascular endothelial cells *in vivo* and *in vitro*. Additionally, *mK5* has a stronger effect on the induction of HUVECs apoptosis [13]. Previous studies have shown that OV-mediated *mK5* can inhibit the growth of colorectal cancer [14] and prostate cancer [15]. Cancer often involves changes in the growth and proliferation of cells. DNA damage and excessive production of reactive oxygen species (ROS) have also been reported to be responsible for the development of cancer [16]. Antioxidant protein containing manganese oxide dismutase (*MnSOD*) is one of the key enzymes for the scavenging of mitochondrial reactive oxygen species (ROS) [17], and is also a novel tumor suppressor protein [18]. Significant growth inhibition was observed in human esophageal squamous [19], PC-3 human prostate cancer [20, 21], MeWo melanoma [22], osteosarcoma [23], ovarian cancer [24,25], MCF-7 breast cancer [26], colorectal cancer [27, 28], hepatocellular carcinoma [29], and pancreatic cancer cell lines [30]. Overexpression of mK5 and MnSOD cDNA by plasmid transfection or adenovirus transduction in various types of cancer triggers growth inhibition *in vivo* and *in vitro*. In this study, *Survivin* promoter was used to regulate the expression of oncolytic adenovirus E1A. The tumor targeting gene viruses Ad-Surp-*mK5* and Ad-Surp-*MnSOD* were constructed for the first time, carrying therapeutic genes *mK5* and *MnSOD* respectively. Moreover, the combined application of two recombinant oncolytic adenoviruses in gastric cancer was studied for the first time. Under both *in vivo* and *in vitro* conditions, the combination of Ad-Surp-*mK5* and Ad-Surp-*MnSOD* was more effective in the inhibition of tumor growth than Ad-Surp-*mK5* or Ad-Surp-*MnSOD* alone. The combination of two recombinant adenoviruses promoted the apoptosis of tumor cells and exhibited no toxic effects on the normal human somatic cells.

## Materials and methods

### Cells and culture

The human gastric cancer cell lines (HGC-27, NCI-87, AGS). Human embryonic kidney cell line (HEK293), human normal lung epithelial cell (Beas-2B), human liver cell (QSG-7701), human colon epithelial cell (NCM-460), and other human cancer cells (HCC-827, HepG-2, Huh-7, A549, SW480, SW620, MDA-MB-231, HeLa, PANC-1, 22RV1, BxPC-3, DU-145, MCF-7, SKOV-3, LNCaP) were purchased from Shanghai Cell Library, Chinese Academy of Sciences (Shanghai, China).

The AGS, HCC-827, HepG-2, SW480, BxPC-3, MCF-7, SKOV-3 and LNCaP were cultured in RPMI-1640 medium, A549 in F-12K medium, MDA-MB-231 in L-15 medium, DU-145 in MEM medium, and other cell lines were cultured in DMEM complete medium. The media was supplemented with 10% fetal bovine serum (Gibco BRL, Grand Island, NY, USA), and all cell lines were incubated in moist air at 37 °C containing 5% CO_2._

### Molecular construction and identification

Adeasy system was used to recombine adenovirus in our laboratory. In this study, the vector of Ad5 virus, shuttle plasmid pShuttle, pCA13, and pCA13-*survivin* were used. The oncolytic adenovirus plasmid pShuttle-E1A-△ E1B with E1B (19 kDa, 55kDa) gene deleted by site directed mutation was previously obtained and preserved, and the *mK5* and *MnSOD* genes were amplified from pXYA05-*mK5* and pXYA05-*MnSOD* respectively by reverse transcription polymerase chain reaction (PCR). The target gene was cloned into pCA13 (by HindIII/XhoI double digestion) in a seamless cloning way to obtain pCA13-gene. The pCA13-*Survivin*, pCA13-*mK5* and pCA13-*MnSOD* were amplified by PCR. The expression box containing the target gene was seamlessly cloned into the corresponding position of the modified shuttle plasmid pShuttle-E1A-△E1B to construct pShuttle-*Surp* (XhoI) -E1A-△ E1B-transgene (BglII). All plasmids were verified by restriction enzyme digestion, PCR and DNA sequence. HEK293 cells were transfected with the Effectene® Transfection Reagent Transfection kit. To amplify the recombinant adenovirus PCR was performed, and the PCR primer sequences and seamless clone primer sequences used in the study are listed in Table 1.

**Table 1.**
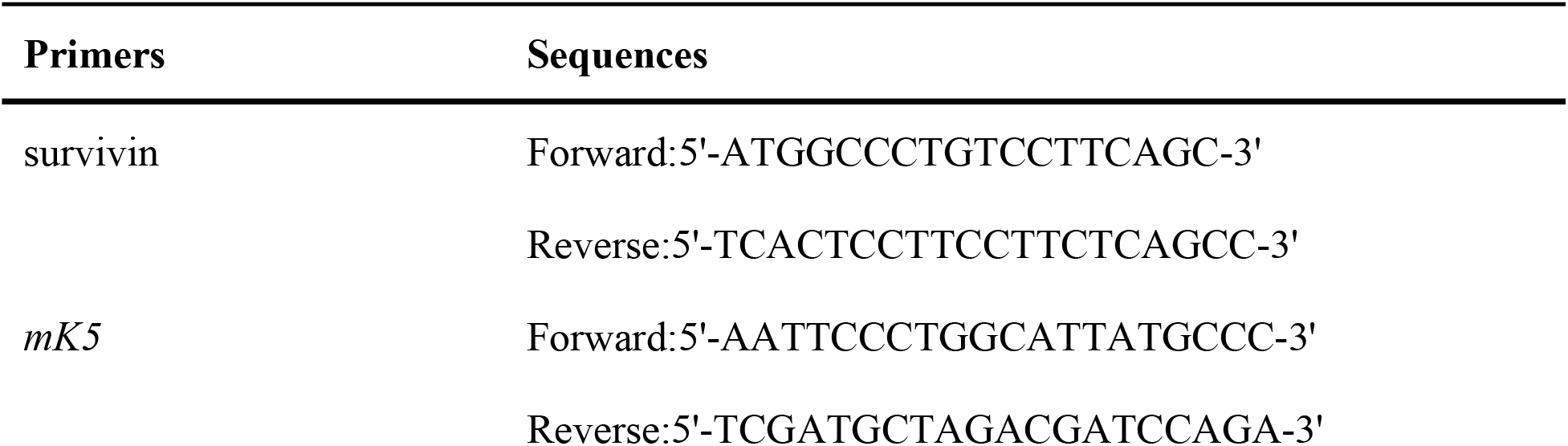
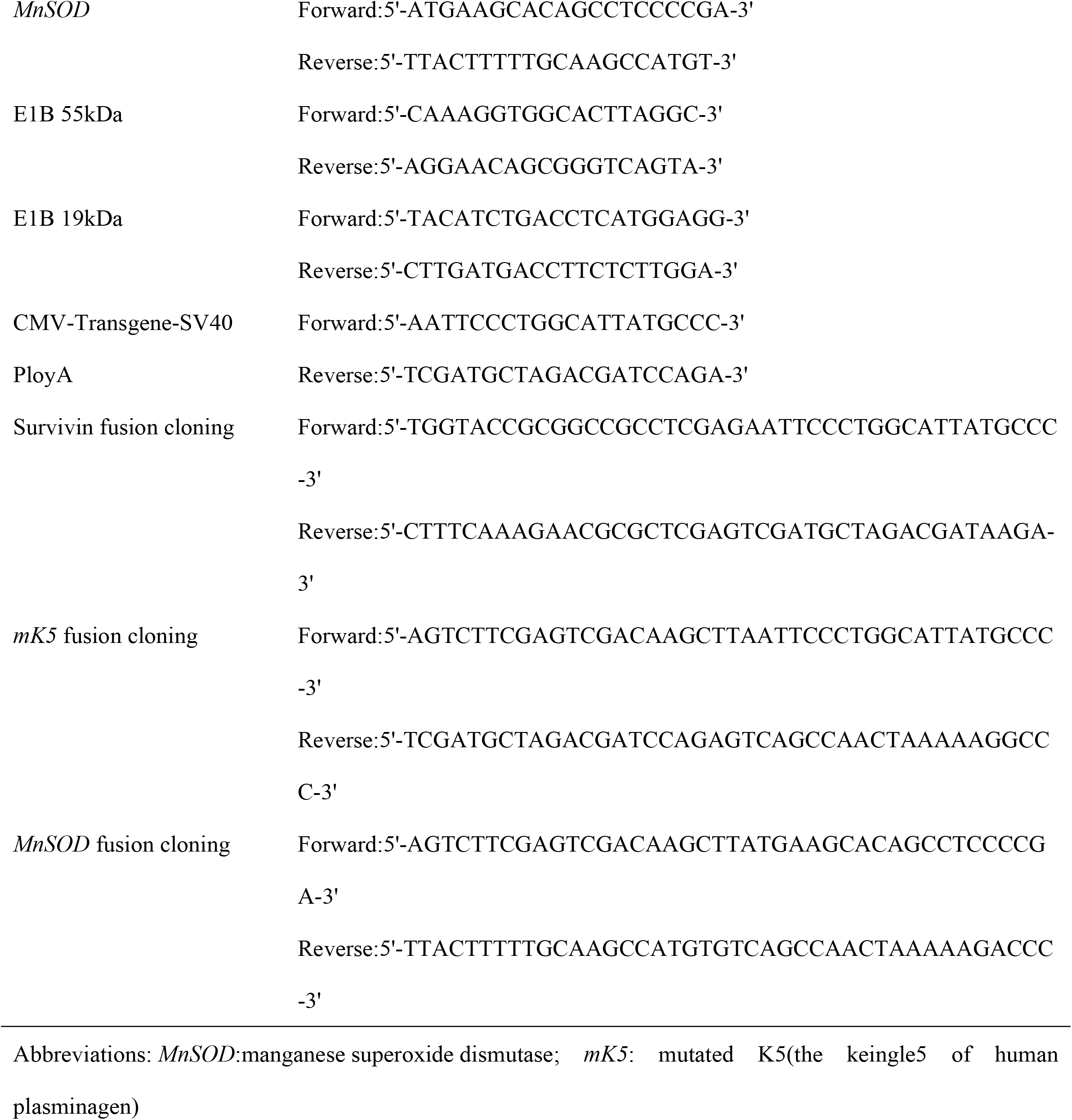
Sequences of primers.

### Western Blot analysis

When multiplicity of infection (MOI) of the recombinant oncolytic adenovirus was 5, GC cells were infected, and the cell lysates were treated and collected after 48 h of incubation. Bradford method was used to detect protein concentration. The samples were run on the 10% polyacrylamide gel electrophoresis, and the proteins were transferred to the polyvinylidene difluoride (PVDF) membrane which was subjected to blocking with 5% skimmed milk powder at room temperature for 30 minutes. This was followed by the incubation of the membranes with primary antibodies at 4 °C for 3 h. The primary antibodies used in the study were Adenovirus-2/5 E1A (1:10000), MnSOD (1:10000), Caspase-3 (1:10000), Caspase-9 (1:10000), Cleaved-PARP (1:10000) (purchased from Santa Cruz Biotechnology. CA. USA), Caspase-8 (1:1000), Bcl-2 (1:1000), Bax (1:1000), SOD2 (1:1000) (purchased from Cell signaling Technology, Beverly, USA), β-actin (1:100000) (purchased from ABclonal Technology, Shanghai, China), *mK5* (1:1000) (purchased from HUABio. Hangzhou, China). This was followed by incubation with the corresponding secondary antibodies for 1.5 h and finally the membranes were scanned for the protein of interest.

### Cell Counting Kit-8 assays

The cells were cultured in 96 well plates at a density of 3000 cells/well for 6 h. To each well 10 µL of the recombinant adenovirus after treatment (Different MOI gradient). Afterwards, CCK-8 detection kit (Beyotime, Shanghai, China) was used to detect cell vitality. The cells were continuously cultured at 37 °C for 4 h, and the absorbance was measured at 450 nm by enzyme-linked immunosorbent assay.

### Flow Cytometry analysis

The GC cells were cultured in 6-well plates and treated with recombinant oncolytic adenovirus for 48 h (MOI = 5). After different treatments, the cells were subjected to treatment with trypsin and collected. Around 100,000 cells were resuspend into 500 µL binding buffer. The analysis was repeated three times according to the manufacturer’s instructions of Annexin V-FITC/PI kit (BD ISRI Sorp, Franklin Lakes, USA).

### EdU Cell proliferation Kit

The GC cells were cultured in 24-well plates were treated with various recombinant oncolytic adenovirus for 48 h (MOI = 5) and then subjected to EdU labeling according to the manufacturer’s instructions for EdU cell proliferation detection kit (Sangon Biotech, Shanghai, China).

### Animal experiment

All animal protocols were approved by the Institutional Animal Handling and Use Committee (Scheme No.: SIBCB-S581-1609-027-C1). The animal welfare and laboratory procedures were strictly carried out in accordance with the Guide to Laboratory Animal Regulations and Standards. Female BALB/c (4-5 weeks) were purchased from Shanghai Experimental Animal Center, Chinese Academy of Sciences (SLAC, Shanghai, China). These experiments were conducted in specific pathogen free (SPF) level laboratory, Shanghai Institute for Biological Sciences, Chinese Academy of Sciences. The HGC-27 cells (1×10^6^) were injected subcutaneously into nude mice to establish the tumor xenograft model. When the tumors were 80-100 mm^3^, the nude mice were divided into 5 groups with five mice in each group. The mice were injected with different recombinant oncolytic adenovirus (1×10^8^ PFU/mouse) or PBS every other day for three consecutive times. Tumor Length and Width was measured every other day after injection to calculate tumor volume:

Volume = (Width^2^ × Length)/2. Finally, tumors were harvested for analysis and the nick terminal labeling of dUTP biotin mediated by terminal deoxynucleotide transferase was performed.

### TUNEL assay

The tumor tissues were fixed with 4% polymethanol for 6 h and subsequently dehydrated with 30% sucrose. The 6 µm tissue sections were analysed by using apoptosis detection kit (red fluorescence, Beyotime). The procedure was carried out according to the manufacturer’s instructions. The specimens were incubated with TUNEL at 37°Cfor 60 min and subjected to staining with 4′,6-diamidino-2-phenylindole (DAPI) at room temperature for 30 min. The specimens were sealed with anti-fluorescence quenching solution and observed under a fluorescence microscope.

### Statistical analysis

The experiments were performed in triplicate and the values are presented as mean ± SD. Student’s t-test or one-way analysis of variance using were used for statistical analysis with the help of Prism GraphPad 8 software. *P < 0.05, **P < 0.01 and ***P < 0.001 were taken as the measure the statistically significant differences.

## Result

### Construction and Characterization of oncolytic adenovirus

To identify the oncolytic adenovirus we had constructed, the corresponding PCR primers were designed and determined its location (Fig 1 A). The target gene was identified by PCR, *mK5*-F primers and *mK5*-R primers were used to identify *mK5* gene (Fig 1 B, channel 4), *MnSOD*-F and *MnSOD*-R primers were used to identify *MnSOD* gene (Fig 1 B, channel 6), *Surp*-F and *Surp*-R primers were used to identify *Survivin* genes (Fig 1B, channel 2), wild-type primers wild-F and wild-R were used to identify wild-type oncolytic adenovirus (Fig 1 B, channels 1, 3 and 5). The gel electrophoresis results showed that the structure of our recombinant oncolytic adenovirus was reasonable. The constructed oncolytic adenovirus was used to infect HGC-27 cells and the protein expression of *mK5* and *MnSOD* were examined by western blot (Fig 1 C), which indicated that the constructed oncolytic adenoviruses could be expressed in gastric cancer cells.

**Fig 1.**
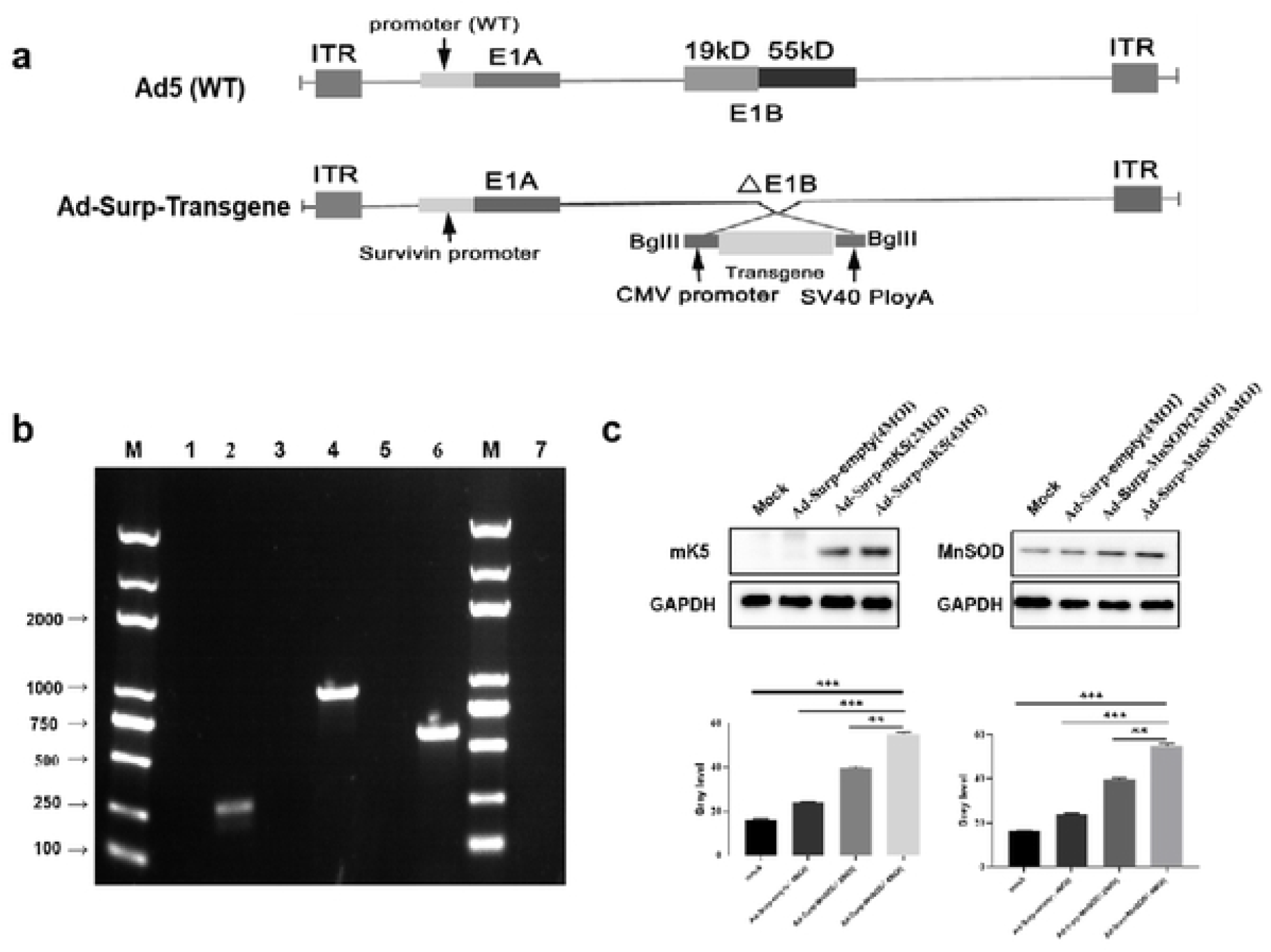
construction and identification of Ad-Surp-Transgene. (a) Transgene includes mutants of *mK5, MnSOD* with wild-type adenovirus Ad5 as skeleton, *mK5:* human plasminogen Kringle5; *MnSOD:* manganese superoxide dismutase. (b) Correlative identification of three kinds of recombinant oncolytic adenovirus Ad-Surp-empty, *Ad-*Surp-*MnSOD* and Ad-Surp-*mK5*. Channels 7 were negative control, and channels 1,3 and 5 were wild virus identification. The *survivin, MnSOD* and *mK5* genes were identified in channels 2, 4 and 6. (c) Western blot analysis showed the protein expression level of *mK5* or SOD-2 in HGC-27 cells treated with Ad-Surp-empty, Ad-Surp-*mK5* or Ad-Surp-*MnSOD*, PBS for 48 hours. Data are shown as the mean ± SD of three independet experiments. * P < 0.05, ** P < 0.01, ***P <0.00 I.

### Selective killing effect of recombinant oncolytic adenovirus on gastric cancer cells in vitro

To detect the killing effect of oncolytic adenovirus *in vitro*, we used Ad-Surp-empty, Ad-Surp-*mK5*, Ad-Surp-*MnSOD*, Ad-Surp-*mK5* and Ad-Surp-*MnSOD* in combination with gradient MOI to infect GC cell lines HGC-27, NCI-87, AGS and human normal cells, human lung epithelial cell Beas-2B, human liver cell QSG-7701, human colon epithelial cell NCM-460 and CCK-8 assayed was used to detect their cell viability for 4 consecutive days. The cell viability of cancer cells treated with various oncolytic adenoviruses decreased significantly (Fig 2 A-F), while the cell activity of all kinds of normal human cells did not decrease significantly. The survival rate of cells at 72 and 96 h was above 80%, indicating that the newly constructed oncolytic adenoviruses had certain selectivity in killing the cells(Fig 2 G-I). At the same time, combination of Ad-Surp-mK5/MnSOD had the strongest killing effect, and the one with significant effect was observed on HGC-27 cells. The proliferation of HGC-27 cells was analyzed by EdU assay and same result was obtained (Fig 3).

**Fig 2.**
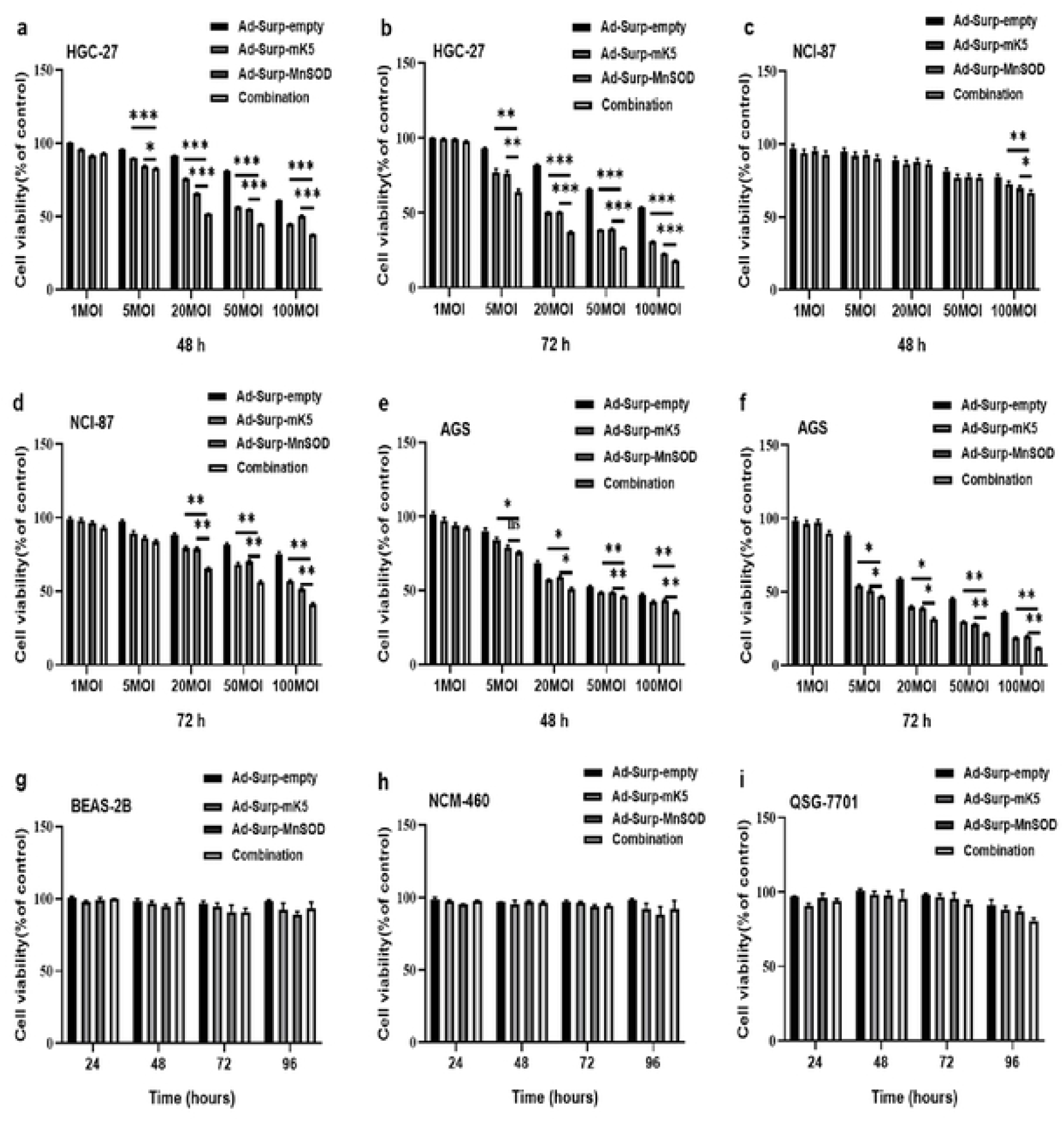
Selective killing effect of recombinant oncolytic adenovirus on gastric cancer. CCK-8 was used to analyze the cell activity of Ad-Surp-Empty, Ad-Surp*-mK5*, Ad-Surp-*MnSOD*, Ad-Surp-*mK5* and Ad-Surp-*MnSOD* combined treatment on HGC-27 cells infecte-d with different viruses for 48h (a) and 72h (b).The cell viability of NCI-87 cells for 48h (c) and 72h (d), AGS cells for 48h (e) and 72h (f) were similarly observed. CCK-8 assay was used to detect the recombinant oncolytic adenovirus alone, Ad-Surp-*mK5* and Ad-Surp-*MnSOD* (1: 1) combined use had no statistical significance on the cell viability of normal human cells Beas-2B (g) NCM-460 (h) and QSG-7701 (i) cells at 24, 48, 72 and 96h when MOI was 20. The data were shown as mean±SD of three independent experiments. * P < 0.05, **P < 0.01, ***P < 0.001.

**Fig 3.**
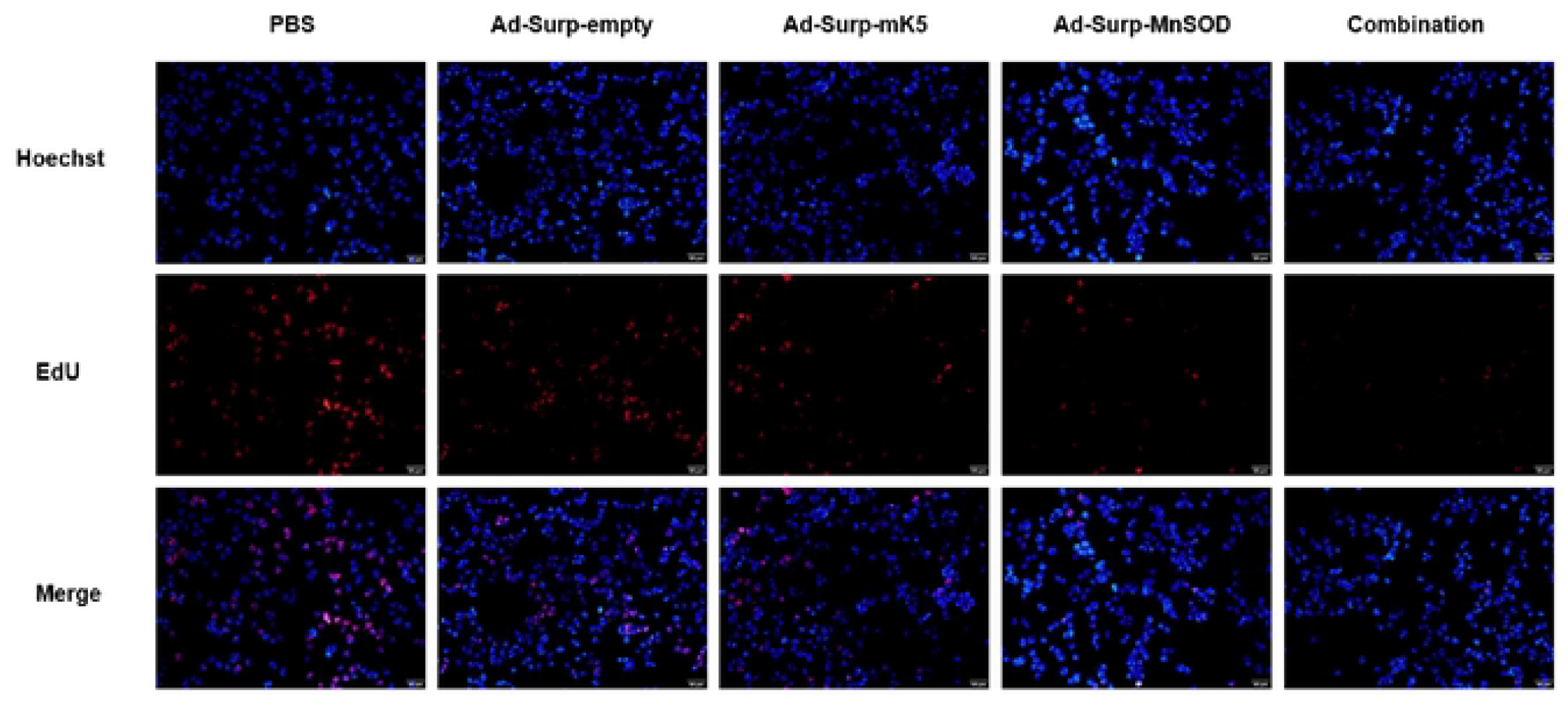
EdU cell proliferation detection. The cell proliferation of Ad-Surp-empty, Ad-Surp-*mK5*, Ad-Surp-*MnSOD*, Ad-Surp-*MnSOD* and Ad-Surp-*mK5* (1:1) combination infected with HGC-27 48 h was observed under fluorescence microscope (MOI=5, Blue fluorescence, Hoechst33342 staining ; Red fluorescence, Apollo staining, scale bar =20 um). The data were shown as mean±SD of three independent experirnents. * P <0.05, **P <0.01, ***P <0.001.

### Apoptosis induced by recombinant oncolytic adenovirus in gastric cancer cells

Next, the recombinant oncolytic adenovirus induced cell death through apoptosis was analysed. We first detected the morphological changes of apoptosis in HGC-27 and QSG-7701 using Hoechst method. The results of the Hoechst 33342 fluorescent staining showed that the apoptotic rate of HGC-27 cells was higher under Ad-Surp-*mK5/MnSOD* (1:1) combination as compared to Ad-Surp-*mK5* or Ad-Surp-*MnSOD* treatments alone (Fig 4 A-E). QSG-7701 cells treated in the same way showed no significant difference in cell morphology compared with the control group (Fig 4 F-J). The apoptotic characteristics were reduced cell volume, condensed nucleoplasm, degradation of the nucleus into fragments and formation of apoptotic bodies. Similarly, flow cytometry analyzer was used to detect the apoptosis of HGC-27 cells (Fig 4L). Ad-Surp-Empty, Ad-Surp-*mK5*, Ad-Surp-*MnSOD*, Ad-Surp-*mK5/MnSOD* (1:1) combined application was used to infect cells for 48 (MOI = 5) hours, and the results showed that the apoptosis of cells in the newly constructed virus combined application group was significantly higher than that in the other groups.

**Fig 4.**
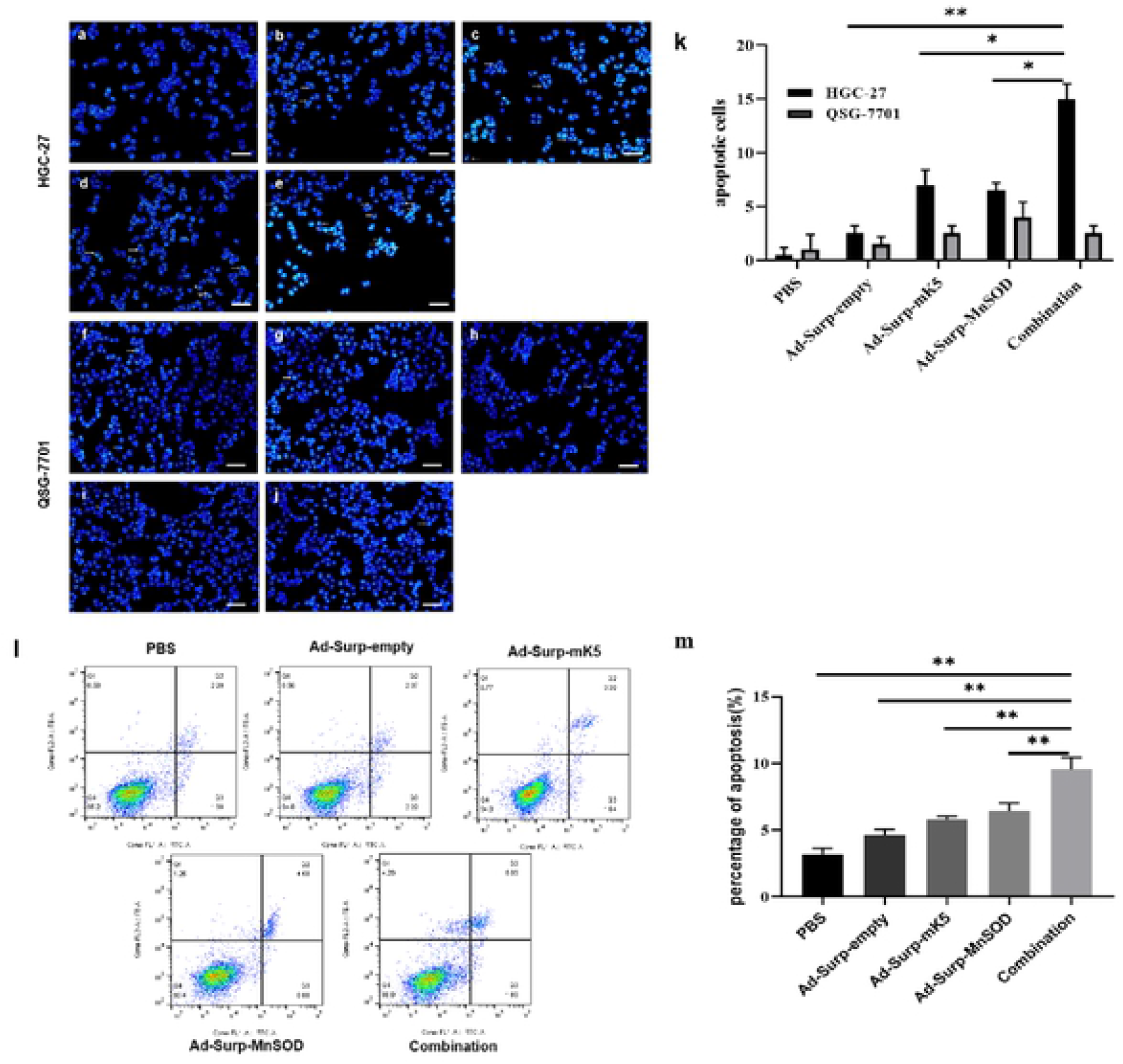
Various recombinant oncolytic adenoviruses selectively induce apoptosis and the mechanism of action. (a-j) HGC-27 (upper two rows) and QSG-7701 (lower two rows) cells were infected with (a,f) phosphate (PBS) (b,g) Ad-Surp-empty (c,h) Ad-Surp-*mK5* (d,i) Ad-Surp-*MnSOD* (e,j) Ad-Surp-*mK5* and Ad-Surp-*MnSOD* ombination (1:1) for 48 h (MOI=5), cell apoptosis was detected by Hoechst33342 staining. Chromatin karyokinization (indicated by arrow) was observed under fluorescence microscope, scale bar=20 um. (k) In figure a-j, HGC-27 and QSG-7701 cells are stained by Hoechst 33342, and the number of apoptotic cells is observed under fluorescence microscopy. (l-m) HGC-27 cells were treated with different recombinant oncolytic adenovirus 48 h, and the PBS treatment group was the control group (MOI=5). The data were shown as mean±SD of three independent experiments. * P < 0.05, ** P < 0.01, *** P < 0.001.

The endogenous mitochondrial apoptotic pathway is the main pathway of mammalian programmed cell death. The oncolytic viruses mainly activate the caspase-dependent apoptosis signaling pathway. Western blotting assay was used to examine the apoptosis related proteins. The results showed that Ad-Surp-*mK5*/*MnSOD* (1:1) combination increased expression of Bax and cleaved-PARP while as the expression of PARP, caspase-3, caspase-9, caspase-8 and Bcl-2 decreased considerably (Fig 5). The effects of Ad-Surp-*mK5*/*MnSOD* (1:1) combination on the apoptosis related proteins were comparatively more significant than Ad-Surp-*mK5* or Ad-Surp-*MnSOD* treatments alone. These results indicate that the Ad-Surp-*mK5*/*MnSOD* (1:1) combination activates the caspase-induced cell signaling pathway in gastric cancer cells more than the other treatment groups and eventually promotes cell apoptosis.

**Fig 5.**
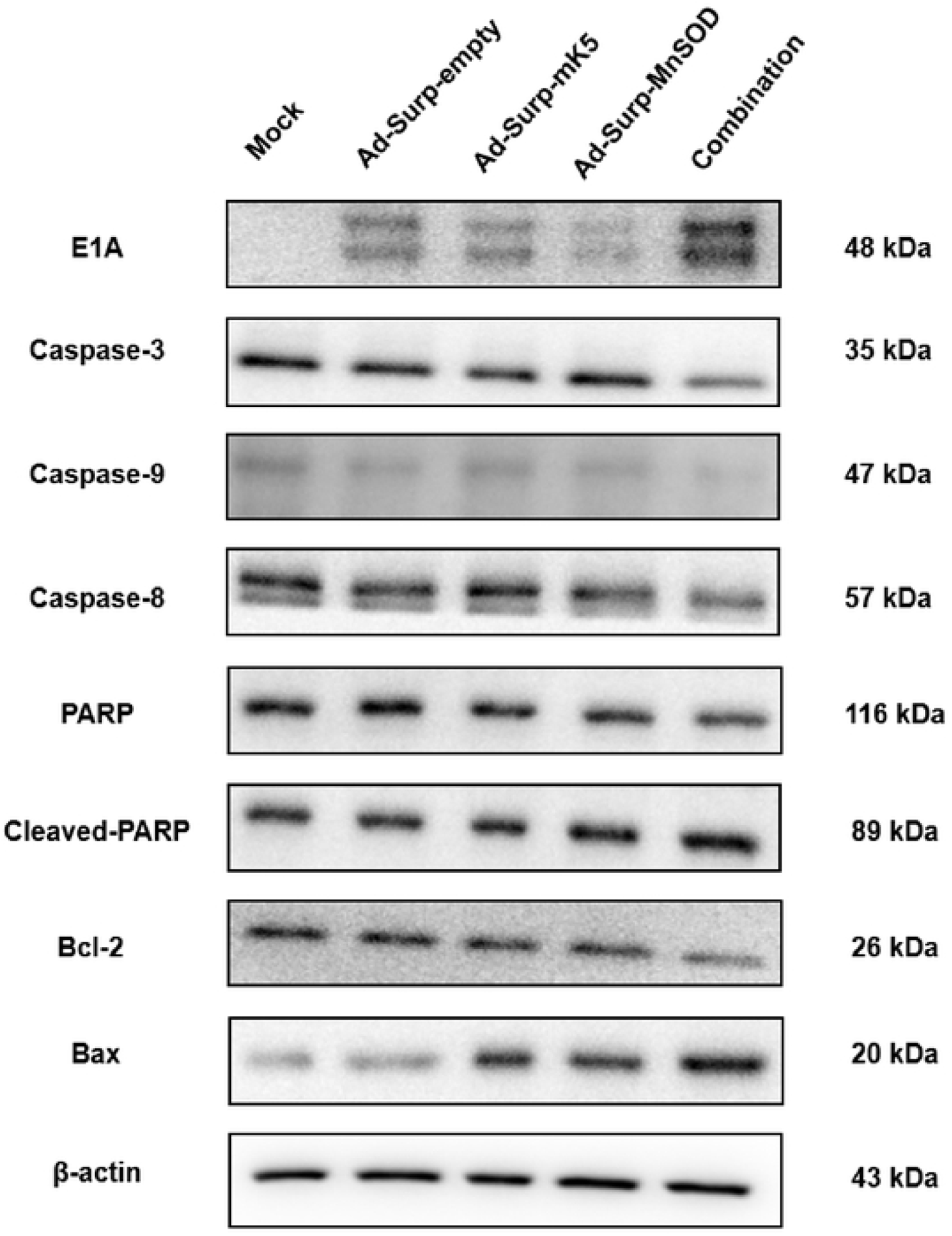
Detection of protein expressions associated with virus proliferation and apoptosisby western blot. As described above, expression of caspase-3, caspase-9, caspase-8, PARP, cleaved-PARP, apoptotic proteins Bcl-2, Bax and virus proliferation-associated protein E1A was observed after treatment with HGC-27 by different viruses.

### Antitumor effect of a newly constructed oncolytic adenovirus Ad-Surp-*mK5/MnSOD* combination *in vivo*

To verify the antitumor activity of the newly constructed oncolytic adenovirus against gastric cancer in nude mice. We established a mouse model of gastric cancer xenotransplantation. The tumor measurement method, drug administration method and final treatment effect on the mice are shown in Fig 6A-B. Compared to the recombinant oncolytic adenovirus alone treatment group and the PBS group, the anti-tumor effect of Ad-Surp-*mK5*/*MnSOD* combination group was significantly enhanced. Additionally, similar therapeutic effect was observed on the tumor growth (Fig 6 C), showing that the tumor volume in the Ad-Surp-*mK5/MnSOD* combination group decreased by 70% compared to the PBS group, and the efficacy was almost double that of the Ad-Surp-*mK5* or Ad-Surp-*MnSOD* alone groups. The body weight of the experimental mice was measured during the experiment (Fig 6 D), there was no significant difference in body weight between the experimental mice in each group and the normal tumor-free nude mice. After tumor resection, the body weight and tumor mass of mice in each group were compared respectively (Fig 6 E-F). There was no significant difference in the body weight of each experimental mouse. However, in the tumor mass analysis of each group, the tumor mass of the virus combination group was significantly lower than those of the other treatment groups. When cells enter apoptosis, DNA ladders are generated by activation of some endonuclease, the exposed 3′-OH is labeled by dUTP catalyzed by terminal deoxynucleic acid transferase (TdT). To further verify the apoptotic status of oncolytic adenovirus infected tumors, the tissue immunofluorescence test was performed (Fig 7). The results showed that the Ad-Surp-*mK5*/*MnSOD* combination treatment group significantly inhibited the tumor growth and induced a higher level of apoptosis as compared to other groups.

**Fig. 6.**
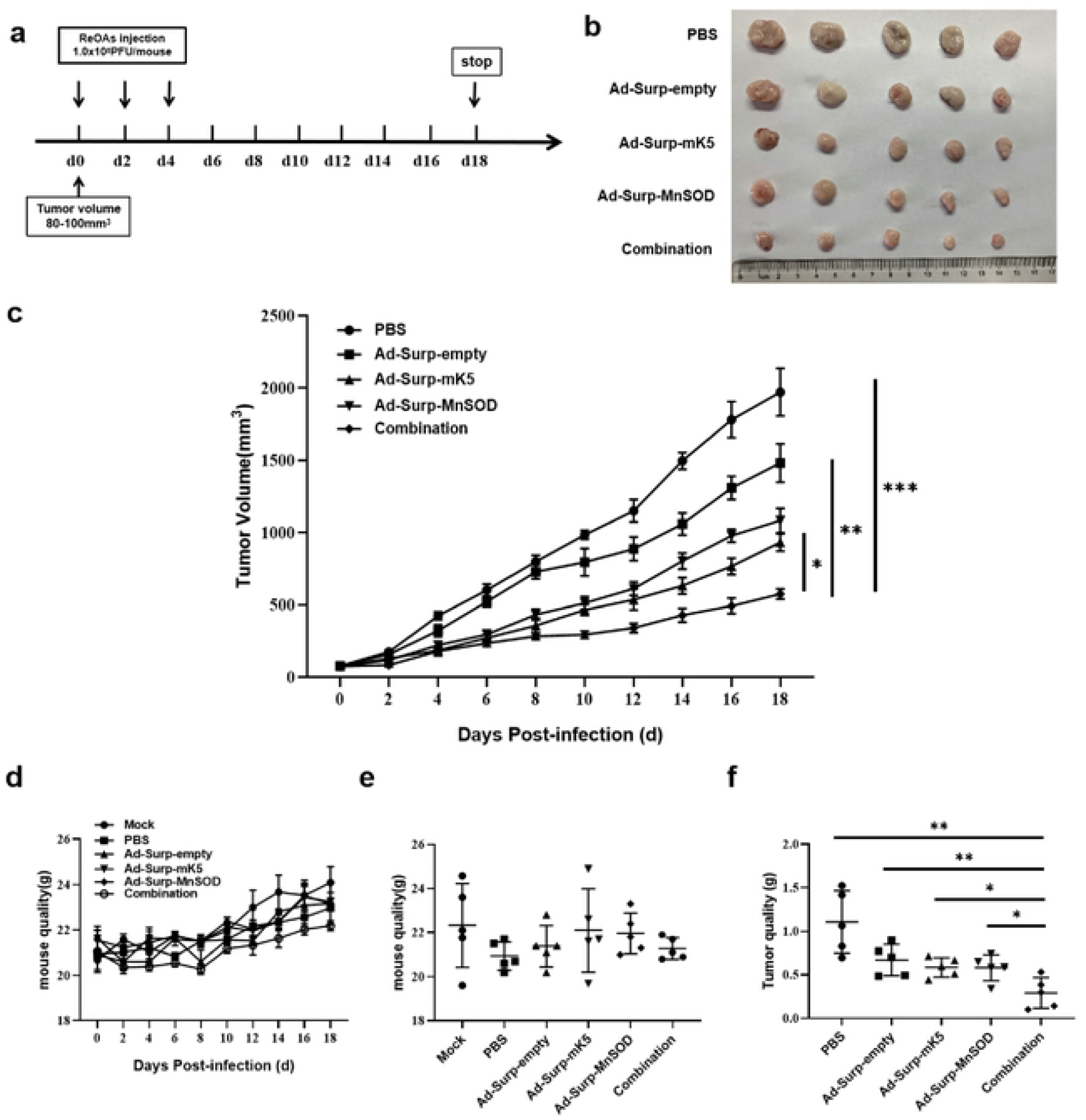
In vivo antitumor effect of recombinant oncolytic adenoviru. HGC-27 cells (1×10^6^) were subcutaneously injected into female BALB/c nude mice to establish a mouse tumor xenograft models. (a) When the tumor grew to 80-100mm^3^, the nude mice were randomly divided into 5 groups (n= 5). The mice were injected with different adenovirus 1.0× 10^8^ PFU/mouse every other day for 3 consecutive intratumor injections. Tumor volume was measured with a vernier caliper every other day, and the experiment was stopped on the 18 th day. (b) Photographs of anatomical tumors of nude mice treated with different adenoviruses. (c) Tumor growth trend of mice Tumor: Volume (mm^3^)= (Length×Width^2^)/2. (d) Changes in body weight of mice in each group during the experiment showed no statistical significance among each group. (e) The final body weight of nude mice was measured by tumor resection after the end of the experiment, and there was no statistical significance among all groups. (f) The final mass of tumors in each group at the end of the experiment. The data were shown as mean±SD of three independent experiments. * P < 0.05, **P < 0.01, ***P < 0.001.

**Fig 7.**
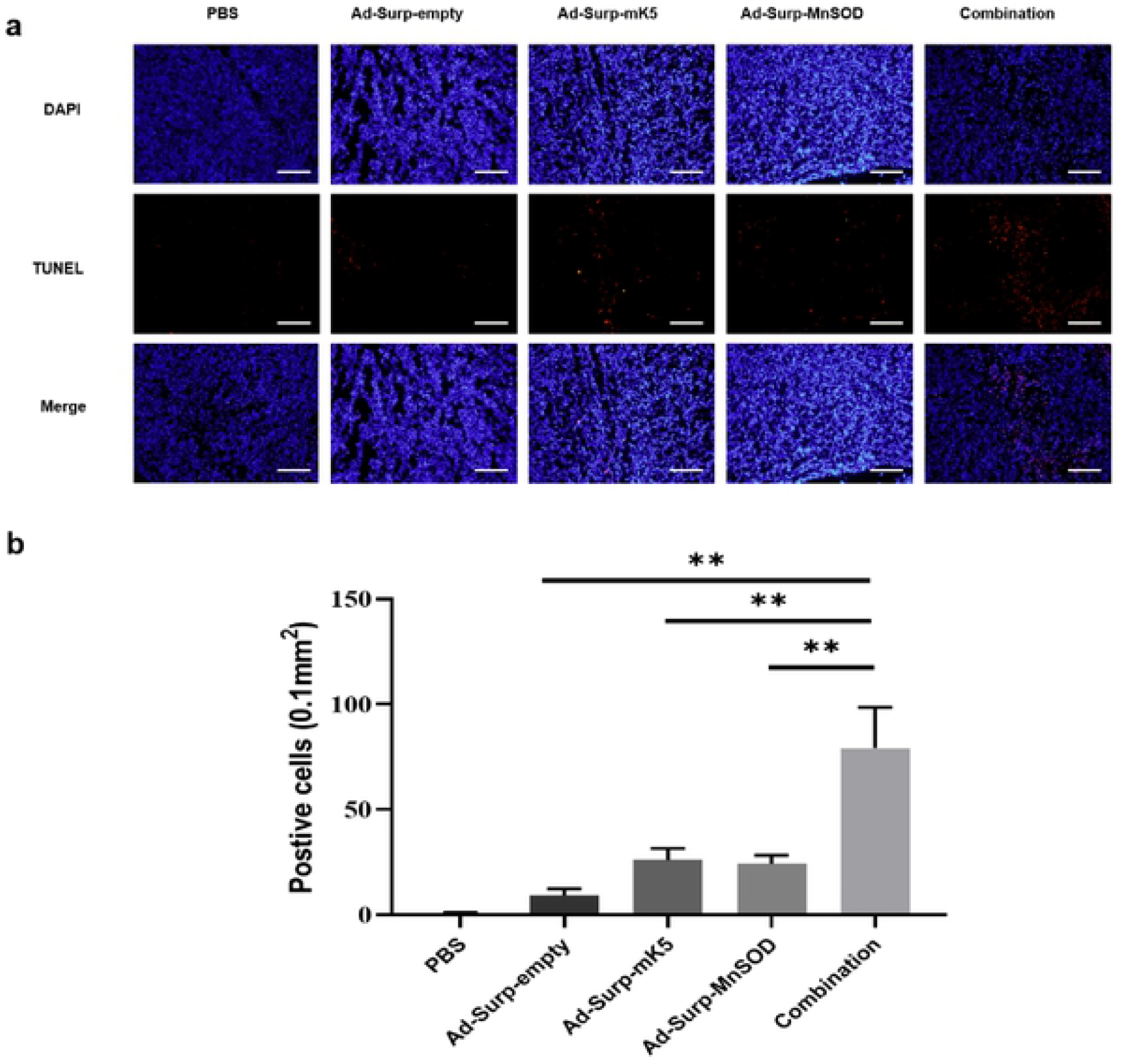
Detection of apoptosis rate of tissue cells by TUNEL staining. (a) Tumor fixation, embedding and cryopreservation were performed after tumor dissection in nude mice, and the thickness of frozen tissue sections was 6 nm (blue fluorescence, DAPI staining; Red fluorescence, TUNEL positive staining), scale bar-200 um. (b) Statistical analysis of positive cells detected by TUNEL staining. The data were shown as mean±SD of three independent experiments. * P <0.05, ** P <0.01, *** P <0.001.

## Discussion

The present study employed a novel strategy called CTGVT [31], to design a dual-regulatory oncolytic adenovirus vector, Ad-Surp-E1A-△ E1B, that can replicate normally and cause tumor cell lysis in survivin-positive GC cells that lack P53 function [32]. Our results indicate that mutant K5 and a novel tumor suppressor gene *MnSOD* can inhibit tumor growth by inducing tumor cell apoptosis. Ad-Surp-*mK5* in combination with Ad-Surp-*MnSOD* (ratio: 1:1) can significantly inhibit the tumor growth in HGC-27 xenograft tumors, and the inhibition effect is more severe than that of the other two recombinant oncolytic adenoviruses alone. These findings suggest that the combined use of Ad-Surp-*mK5* and Ad-Surp-*MnSOD* has great potential in the treatment of GC. Li et al. showed that *mK5* can induce apoptosis of endothelial cells both *in vivo* and *in vitro*, and is a more effective anti-angiogenic drug than K5 [13]. At the same time, in mature endothelial cells, ROS is an important mediator of different signaling pathways related to angiogenesis and cell metabolism, regulation of cell proliferation, migration and gene expression [33]. Hypoxia-induced vascular endothelial growth factor (VEGF) expression plays a key role in the promotion of tumor angiogenesis, however, the molecular mechanism underlying the regulation of VEGF expression under hypoxia remains unclear. Xu et al. found that induction of VEGF in hypoxic HepG2 cells was ROS-dependent [34]. Studies have shown that ROS-mediated signals play a key role in the regulation of transcription during hypoxia, and it has been reported that ROS-dependent mechanisms induce VEGF to enter cancer cells under hypoxia conditions. Existing ROS scavengers can reduce the expression of VEGF [35], while *MnSOD* is the main antioxidant in mitochondria, which can resist ROS by catalyzing the transformation of ROS from superoxide anion (O^2-^) to hydrogen peroxide (H_2_O_2_) in mitochondria [36]. The deletion of *MnSOD* in various tumors suggests that *MnSOD* may be a candidate tumor suppressor gene.

In the present study, we constructed two novel survivin-regulated, dual-targeted oncolytic adenovirus Ad-Surp-*mK5/MnSOD*, which carries the gene *mK5* or *MnSOD*. Our current data revealed that the anti-tumor efficacy of Ad-Surp-*mK5* and Ad-Surp-*MnSOD* (1:1) combination in the treatment of gastric cancer is significantly better than that of Ad-Surp-*mK5* and Ad-Surp-*MnSOD* alone. It was further demonstrated that compared to the recombinant adenovirus alone, the combination of the virus induced apoptosis of gastric cancer cells specifically and the expression of caspase was stronger. Additionally, no toxicity was found in human lung epithelial cells, human liver cells and human colon cells in the *in vitro* as revealed by CCK-8 assay. Above data show that our new construct is a potential candidate for experimental GC anticancer drug, Using CTGVT strategy to ensure the safety of viral vector we put the virus E1B knockout Ad5 as the initial viral vector, but simply deleted the viral gene does not stop unnecessary tissue damage, tumor selectivity can be enhanced by the placement of a tumor-specific promoter that is transcribed via mRNA upstream of a critical viral gene (E1A), so we used a novel *Survivin* promoter. [37]. *Survivin* promoter regulation of E1A gene makes up for the lack of selectivity of OVs vector. Adenovirus with anticancer therapy gene after recombination proliferates and replicates in gastric cancer cells, triggering the release of progeny virus and which subsequently infect and eventually destroy and kill the cancer cells. Chen et al. reported that when a system-managed suicide gene is driven by a *Survivin* promoter, systemic toxicity may be significantly reduced [38]. Meanwhile, *Survivin* promoters have been shown to induce transgenic overexpression in cancer [11].

Oncolytic therapy is another form of immune stimulation that works through the patient’s own immune system to combat tumor growth and facilitate tumor removal [39]. Although our novel survivin-regulated, dual-targeted OVs have achieved powerful antitumor effects in GC cells and animal models, there are still some challenges that need to be further addressed. The antiproliferative effects of Ad-Surp-*mK5*/*MnSOD* against the GC cells still remains low. Although they carry therapeutic genes *mK5* or/and *MnSOD* and soluble tumor recombinant adenovirus effectively inhibit the growth of nude mice tumors, they can’t completely eradicate tumor xenograft. The primary reason for this result may be the host immune response and the limited distribution of OVs in the tumor tissues [40]. The type and size capsule of viruses and the nature of the tumor microenvironment affects the penetration and diffusion of OVs [41]. Therefore, it is urgent to address such problems in future virus-gene targeted cancer therapy and to achieve more significant therapeutic effects through the combination of other conventional therapies.

In conclusion, OV successfully constructed by CTGVT strategies, through *Survivin* promoter regulation targeting tumor adenovirus vector which makes the recombinant adenovirus in GC cells more specific. It can effectively mediated *mK5* or *MnSOD* gene expression maximizing the role of dissolving the tumor and inhibiting the tumor growth while minimizing side effects on normal tissues. Therefore, it has a wider application and may prove to be an important candidate for the treatment of GC.

## Acknowledgement

The authors would like to thank the staff of Shanghai Institute for Biological Sciences and our colleagues for their support to this study.

